# Large-scale network dynamics of beta-band oscillations underlie auditory perceptual decision-making

**DOI:** 10.1101/095356

**Authors:** Mohsen Alavash, Christoph Daube, Malte Wöestmann, Alex Brandmeyer, Jonas Obleser

## Abstract

Perceptual decisions vary in the speed at which we make them. Evidence suggests that translating sensory information into behavioral decisions relies on distributed interacting neural populations, with decision speed hinging on power modulations of neural oscillations. Yet, the dependence of perceptual decisions on the large-scale network organization of coupled neural oscillations has remained elusive. We measured magnetoencephalography signals in human listeners who judged acoustic stimuli made of carefully titrated clouds of tone sweeps. These stimuli were used under two task contexts where the participants judged the overall pitch or direction of the tone sweeps. We traced the large-scale network dynamics of source-projected neural oscillations on a trial-by-trial basis using power envelope correlations and graph-theoretical network discovery. Under both tasks, faster decisions were predicted by higher segregation and lower integration of coupled beta-band (~16-28 Hz) oscillations. We also uncovered brain network states that promoted faster decisions and emerged from lower-order auditory and higher-order control brain areas. Specifically, decision speed in judging tone-sweep direction critically relied on nodal network configurations of anterior temporal, cingulate and middle frontal cortices. Our findings suggest that global network communication during perceptual decision-making is implemented in the human brain by large-scale couplings between beta-band neural oscillations.

**Author Summary:** The speed at which we make perceptual decisions varies. This translation of sensory information into behavioral decisions hinges on dynamic changes in neural oscillatory activity. However, the large-scale neural network embodiment supporting perceptual decision-making is unclear. Alavash et al. address this question by experimenting two auditory perceptual decision-making situations. Using graph-theoretical network discovery, they trace the large-scale network dynamics of coupled neural oscillations to uncover brain network states supporting the speed of auditory perceptual decisions. They find that higher network segregation of coupled beta-band oscillations supports faster auditory perceptual decisions over trials. Moreover, when auditory perceptual decisions are relatively difficult, the decision speed benefits from higher segregation of frontal cortical areas, but lower segregation and integration of auditory cortical areas.

## Introduction

Contemporary neuroscience is departing from a focus on regional brain activations towards emphasizing the network organization of brain function. This view recognizes the large-scale interactions between distributed cortical areas as the biological basis of behavior and cognition (Sporns, 2014; Misic and Sporns, 2016). A more mechanistic view holds frequency-specific neural oscillations to be relevant to behavior and cognition (Buzsaki and Draguhn, 2004; Engel et al., 2013). How the large-scale network organization of interacting neural oscillations (Palva and Palva, 2012; Siegel et al., 2012), in particular their temporal network dynamics (Kopell et al., 2014; Shine et al., 2016), relate to perception and cognition is poorly understood. Here, we investigate the dependence of auditory perceptual decision-making in humans on spectrally-, temporally- and topologically-resolved large-scale brain networks.

Accumulating evidence suggests that frequency-specific neural oscillations are key to processing sensory information (Palva et al., 2010; Hanslmayr et al., 2011; de Pesters et al., 2016). For example, previous studies indicate that attentional modulation of cortical excitability in sensory regions is reflected in oscillatory alpha power (~8-10 Hz) under visual (Jensen and Mazaheri, 2010; Lange et al., 2013; Lou et al., 2014) or auditory tasks (Müller and Weisz, 2012; Strauß et al., 2014; Weisz et al., 2014a; Wöstmann et al., 2016). Additionally, it has been shown that audiovisual perception relies on synchronized cortical networks within beta (~20 Hz) and gamma (~80 Hz) bands (Hipp et al., 2011). Recently, studies have begun to explore more specifically whether modulations in neural oscillations arise from lower-order sensory or higher-order control areas (Park et al., 2015; Friese et al., 2016; Kayser et al., 2016; Sadaghiani and Kleinschmidt, 2016). Here, based on localization of neurophysiological sources (Hillebrand et al., 2005), we explore the large-scale network organization of interacting neural oscillations during auditory processing.

Specifically, we ask how the network topology of coupled neural oscillations (Bassett et al., 2009) relates to the listeners’ perceptual decisions. In a previous magnetoencephalography (MEG) study, Nicol and colleagues measured synchronization of brain gamma-band (33-64 Hz) responses in an auditory mismatch-negativity paradigm (Nicol et al., 2012). They found that deviant stimuli were associated with increase in local network clustering in left temporal sensors within the immediate response period. Building upon pre-stimulus hemodynamic responses, Sadaghiani et al. (2015) recently suggested higher modularity of brain networks as a proxy for perceiving near-threshold auditory tones. Moreover, it has been shown that higher global integration of brain networks measured from pre-stimulus high-alpha band MEG responses precede the detection of near-threshold stimuli (Weisz et al., 2014b; Leske et al., 2015). In sum, brain network correlates of auditory perception have been observed on different topological scales.

Naturally, cortical networks involved in processing sensory information require context-sensitive configurations, and also moment-to-moment reconfigurations to fulfill dynamic task adjustments (Bassett et al., 2006). This leads the neural co-activations, which shape the brain functional connectivity, to diverge from their underlying structural connectivity (Park and Friston, 2013; Marrelec et al., 2016; Misic et al., 2016). As such, estimation of functional connectivity, when collapsed over time, overemphasizes structural connectivity (Honey et al., 2007; Shen et al., 2015) and disregards the temporal dynamics of large-scale brain network topology (Zalesky et al., 2014; Kringelbach et al., 2015). It is these dynamics that have been found relevant to behavior during challenging motor or cognitive tasks (Bassett et al., 2013; Braun et al., 2015; Alavash et al., 2016; Chai et al., 2016).

Therefore, to find the neural network substrate of auditory perceptual decision-making, we adopt the framework of dynamic brain networks (Calhoun et al., 2014; Deco et al., 2015) and merge this with neural oscillations to uncover frequency-specific brain network states. Our method is based on a previously established technique to estimate large-scale neural interactions in source space (Hipp et al., 2012) and graph-theoretical network analysis (Bullmore and Sporns, 2009).

We apply this approach to MEG signals measured from human listeners who made perceptual decisions on brief acoustic textures under two distinct task sets. The acoustic textures consisted of densely layered tone sweeps which varied in their overall pitch (high or low), as well as the proportion of coherent tones in terms of sweep direction (up or down; Figure 1A). Using the identical set of stimuli, two auditory paradigms with distinct decision contexts were designed to deliver challenging perceptual decision-making tasks (Figure 1B). As such, the individuals’ perceptual decision accuracy and speed fluctuated on a trial-by-trial basis (Figure 1C). This allowed us to investigate the relation between frequency-specific brain network states and trial-by-trial decision-making performance (Figure 2). Since, under each of the two perceptual decision-making tasks, subjects focused on a different acoustic feature of an identical set of auditory stimuli, two dynamic network profiles were expected. First, we anticipated brain network states responsible for the cortico-cortical communication (mainly fronto-temporal) involved in common for both tasks to predict the decision-making performance. Second, we expected task-specific brain network states emerging from auditory association or higher-order decision areas to differentially predict the performance in either of the tasks.

**Figure 1.**
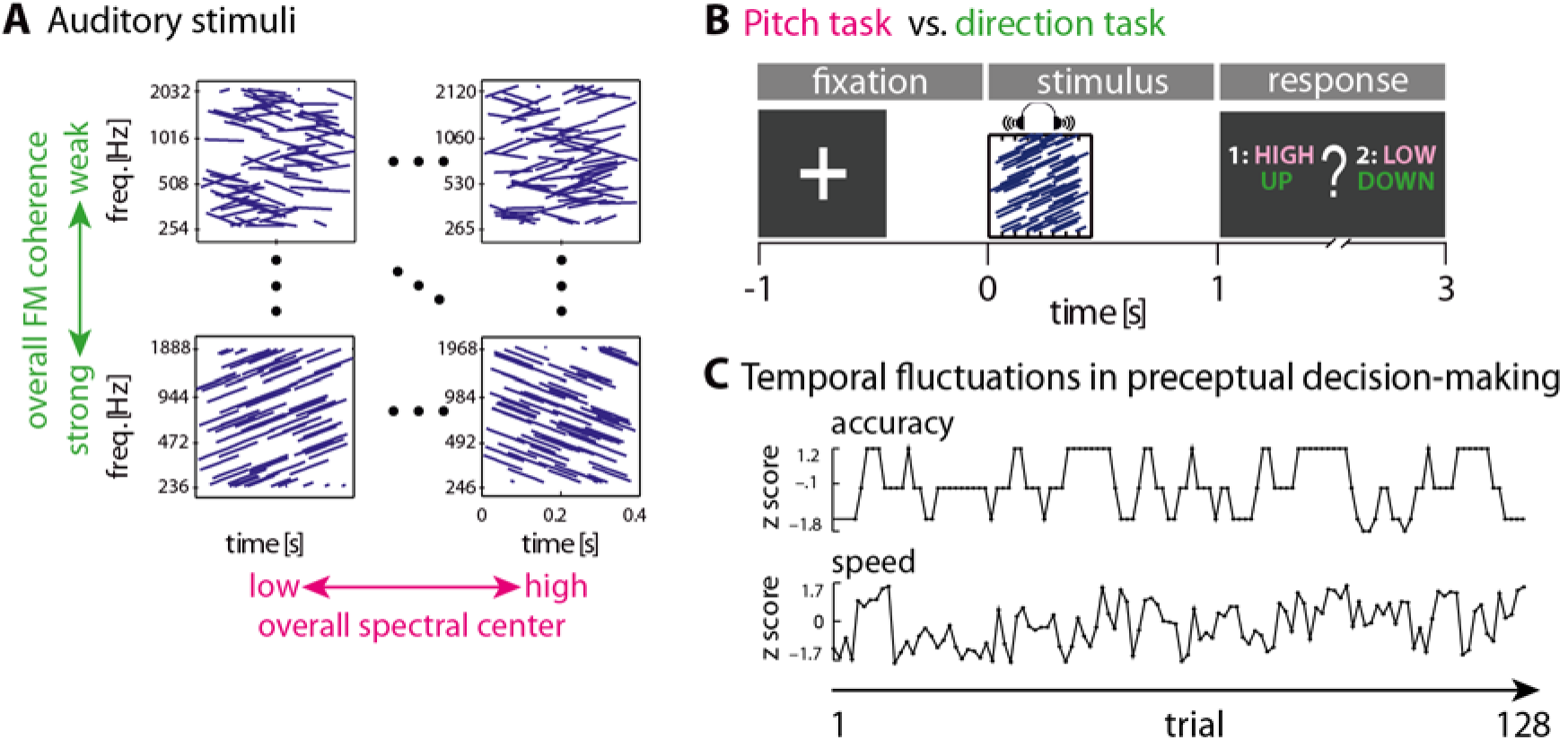
Experimental stimuli and tasks. **(A)** Auditory stimuli used to design the tasks. Each cell represents an acoustic texture which can be viewed as a pattern of sound sweeps whose frequency increases or decreases over time. A texture stimulus had a duration of 400 ms, and consisted of 72 frequency modulated (FM) sweeps of 100 ms duration. The stimuli where titrated along two dimensions: overall coherence and spectral center. For a given stimulus, a variable proportion (25-100%) of the sweeps were assigned the same frequency slope (coherence), i.e. their frequency was going up or down at the same rate over time. The rest of the sweeps had a randomly assigned slope. In addition, each stimulus had one of eight spectral centers relative to a mean center frequency. **(B)** Two auditory perceptual decision-making tasks, namely *pitch* and *direction,* were designed using the identical acoustic stimuli. During the pitch task, the subjects judged the overall pitch of the stimuli (low or high). In the direction task, they were asked to report the overall direction (up or down) in which the frequency of the stimuli was changing (increasing or decreasing) over time. Subjects had 3 seconds at maximum to press one of two response buttons to report their perceptual decision. In each task, the decision labels for the left hand and right hand buttons (indicated by 1/2) were randomized across trials, and were shown after stimulus presentation within the response window. There were eight blocks per task, and the order of the tasks alternated from one block to another. **(C)** Exemplary trial-by-trial auditory perceptual decision accuracy (moving average of four trials applied to correct/incorrect responses) and decision speed ([response time]^−1^). Before the actual tasks, an adaptive perceptual tracking was used to tailor the two tasks per participant, so that their overall accuracy converged at ~70%. This led the individuals’ decision accuracy and speed to fluctuate over trials. Note that for the purpose of the regression analysis (see ‘Materials and Methods’), the trial-wise estimates of accuracy and speed were first rank-transformed and then normalized (i.e., z-scored). Exemplary data are shown for a representative participant; pitch task, second block.

**Figure 2.**
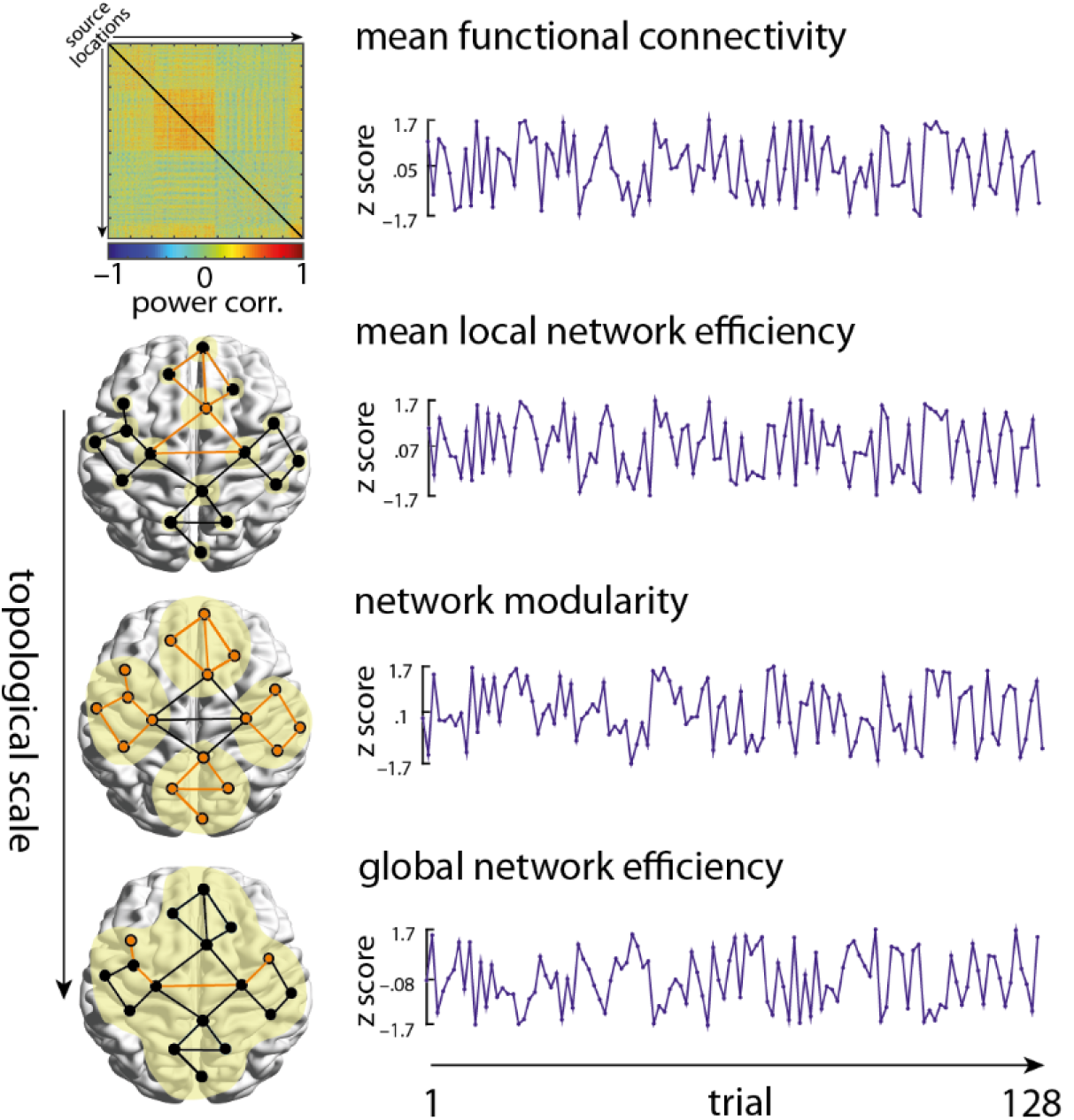
Trial-by-trial dynamics of brain functional connectivity and network topology. To investigate the relation between the frequency-specific brain network states and decision-making performance, all-to-all power envelope correlations (source connectivity matrix) and whole-brain graph-theoretical network metrics were estimated at 10% of network density (see ‘Materials and Methods’). This analysis was done at each frequency within 1-32 Hz, and per trial in the course of each pitch and direction task. The temporal graph-theoretical metrics captured brain network states on the local (local efficiency), intermediate (modularity) and global (global efficiency) scales of network topology. The yellow shaded ovals illustrate the topological scale at which a network metric is measured. *Global efficiency* (bottom graph) is inversely related to the sum of shortest path lengths (e.g. orange path) between every pair of nodes. *Mean local efficiency* (top graph) is equivalent to global efficiency computed on the direct neighbors of each node (e.g. orange node) which is then averaged over all nodes. *Modularity* (middle graph) describes the segregation of partner nodes into relatively dense groups (here, the orange nodes forming four modules) which are sparsely interconnected. For the purpose of the regression analysis (see ‘Materials and Methods’), the trial-wise estimates of network metrics were first rank-transformed and then normalized (i.e., z-scored). Exemplary data are shown for a representative subject as in Figure 1C.

## Results

### Auditory perceptual decision-making performance

The participants judged the overall pitch or sweep direction of the acoustic texture stimuli, and showed accuracies around 70% as it had been intended with the adaptive tracking procedure (average accuracy[%]±SEM: pitch task=76% ±1.1, direction task=70% ±2.1; mean of decision speed [*s*^−1^]±SEM: pitch task=1.8±0.1, direction task=1.7±0.1). The bootstrap Kolmogorov-Smirnov test revealed that the distributions of the behavioral measures did not significantly deviate from a normal distribution (accuracy: pitch task p=0.8; direction task p=0.1; decision speed: pitch task p=0.8; direction task p=0.5). Despite our experimental efforts to equate the difficulty in both tasks, analysis of variance (ANOVA) revealed a main effect of task for both accuracy (F(1,19)=6, *η_G_*^2^, p<0.05) and decision speed (F(1,19)=9.9, *η_G_*^2^, p<0.01). Participants showed significantly lower accuracy in the direction task as compared to the pitch task (exact permutation test for a paired test; p<0.05). In addition, decision speed in the direction task was significantly lower as compared to the pitch task (p<0.01). The experimental manipulation of the pitch of the acoustic textures yielded significant effects of stimulus spectral center on both accuracy (F(3,57)=64, *η_G_*^2^, p<0.001) and decision speed (F(3,57)=30, *η_G_*^2^, p<0.05). However, stimulus coherence had a significant effect only on accuracy (F(3,57)=45.3, *η_G_*^2^ =0.12, p<0.001). Across participants, there was a significant positive correlation between decision speed in the pitch task and decision speed in the direction task (Spearman's *ρ* =0.9; *p*<0.01). However, the correlation between accuracies in the two tasks was not significant (*ρ* =0.2; p=0.47).

The correlation between trial-by-trial estimates of decision accuracy or speed with trial-by-trial acoustic features of the stimuli was tested using a two-level regression analysis (see ‘Materials and Methods’). We found a significant positive correlation between decision speed and stimulus coherence in the direction task (average regression weights±SEM: 0.045±0.012; one-sample exact permutation test: p<0.01). Additionally, trial-by-trial estimates of decision accuracy in the direction task showed a significant positive correlation with the coherence of the stimuli (average regression weights±SEM: 0.087±0.01; p<0.01). There was also a significant negative correlation between decision accuracy in the pitch task and stimulus coherence (average regression weight±SEM: −0.022±0.01; p<0.05).

### Neural oscillatory power during auditory perceptual decision-making

We investigated power perturbations in the MEG oscillatory signal while subjects listened to the acoustic textures, and judged their overall pitch or sweep direction. As it is illustrated in Figure 3, MEG oscillatory alpha (~8-13 Hz) power was increased relative to the baseline interval (-0.5 to 0 s) just after stimulus presentation. In addition, during and after stimulus presentation but before the response prompt (0 to 1s), we observed a left-lateralized decrease in the MEG oscillatory power in the low- and mid-beta band (~14-24 Hz) relative to the baseline interval. The above perturbations in alpha and beta bands were similarly observed in both pitch and direction tasks (Figure 3A and B, first two panels) and are well in line with previous studies on the neural substrates of perceptual decision-making (Donner et al., 2009; Haegens et al., 2011; O'Connell et al., 2012; Kelly and O'Connell, 2015). Finally, as expected, there was a strong, motor-related suppression in the MEG oscillatory power relative to baseline within the time interval when the subjects manually reported their perceptual decision following the response prompt (Pfurtscheller and Lopes da Silva, 1999). This perturbation was widely-distributed within alpha and beta bands (8-32 Hz; Figure 3A and B, first panel).

**Figure 3.**
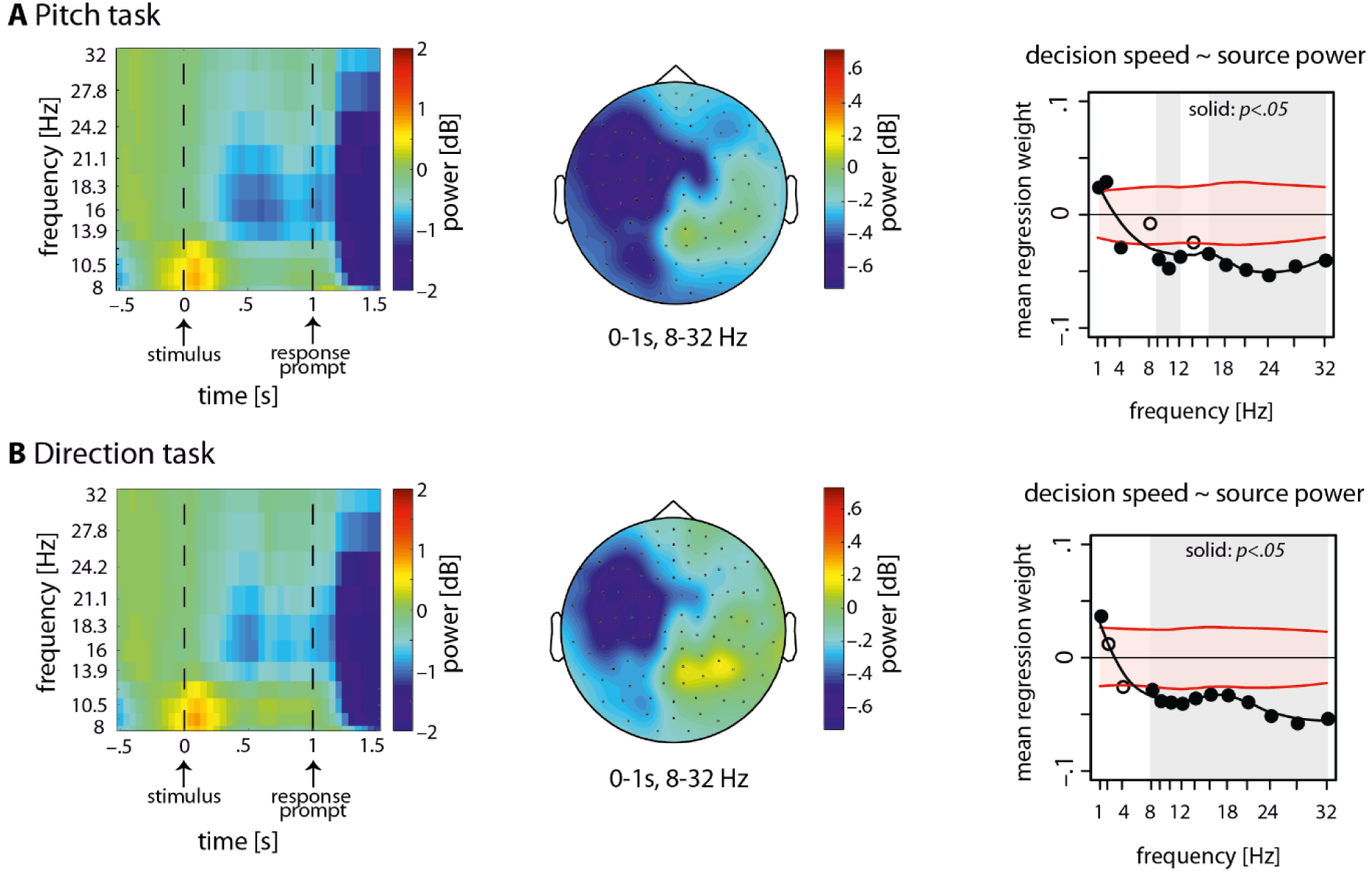
Dynamics of neural oscillatory power under each auditory task and their relation to the speed of perceptual decisions. Spectrotemporal representations of the epoched signals during **(A)** the pitch task and **(B)** the direction task were estimated and averaged over trials, 102 combined gradiometer sensors and all subjects (N=20). While an increase in oscillatory alpha (8-13 Hz) power was observed time-locked to the auditory stimulation relative to the baseline interval (−0.5 to 0s), there was a decrease in the oscillatory power within low- and mid-beta bands (14–24 Hz) during the time window when the participants listened to the stimuli (0 to 1s). The topographical maps show the broad-band baseline-corrected oscillatory power (8–32 Hz) from stimulus onset to the onset of the response prompt (0-1s). Note that within this time period, the subjects were not yet aware of the mapping between the decision labels (pitch task: high/low; direction task: up/down) and left/right hand response buttons, since the mapping was randomized across trials. *Regression analysis.* The relation between the ongoing power of neural oscillations and decision speed was investigated by means of linear regression where trial-by-trial decision speed was predicted by the oscillatory power of source signals. In the course of both auditory tasks, faster perceptual decisions negatively correlated with the ongoing oscillatory power of source signals within alpha and beta bands. *Black circles.* The normalized regression weights averaged over subjects at each frequency. *Horizontal shades.* 95% confidence interval of the null mean regression weights generated by circularly shifting the behavioral responses across trials (corrected for multiple comparisons across frequency bins using false-coverage statement rate (FCR; p<0.05)). For visualization purposes, the smooth curves were estimated using local polynomial regression fitting.

Since the aim of this study was to find the relation between frequency-specific brain network states and auditory perceptual decision-making on a trial-by-trial basis, we next focused on the trial-by-trial fluctuations in the power of the source-projected signals. To this end, we implemented a general linear model (GLM) per participant whereby time series estimates of trial-by-trial decision accuracy or speed (Figure 1C) were predicted by the baseline-corrected power of the whole-brain source-projected signals. We found significant negative correlations between decision speed during pitch or direction task and the ongoing neural oscillatory power within alpha and beta bands (8-32 Hz; Figure 3A and B, third panels). These results indicate that, during both pitch and direction tasks, a stronger decrease in the neural oscillatory power relative to the baseline interval correlated with faster perceptual decisions. However, we found no significant correlation between decision accuracy and the ongoing oscillatory power of source-projected signals.

### Whole-brain network dynamics of beta-band oscillations predict decision speed

The aim of this study was to find frequency-specific brain network states underlying individuals’ perceptual decision-making in the course of judging auditory stimuli. The auditory stimuli were identical, but presented in two distinct task sets, i.e. either judging the overall pitch or the overall direction of frequency-modulated (FM) tone sweeps. To predict trial-by-trial decisionmaking performance from the ongoing brain network states, we implemented a linear regression model where time series estimates of trial-by-trial decision accuracy or speed (Figure 1C) were predicted by the temporal graph-theoretical network metrics (Figure 2).

On the whole-brain level, and for both pitch and direction tasks, we found significant correlations between decision speed on the one hand, and the functional connectivity and the topology of dynamic brain networks on the other hand (Figure 4). These correlations peaked within the beta band range (Figure 4, solid points). The significant correlations indicate that, for both pitch and direction task, higher local efficiency but lower global efficiency of large-scale brain networks supported faster perceptual decisions (Figure 4, second and last columns, respectively). In addition, higher segregation of brain network modules predicted faster perceptual decisions in both tasks (Figure 4, third column).

**Figure 4.**
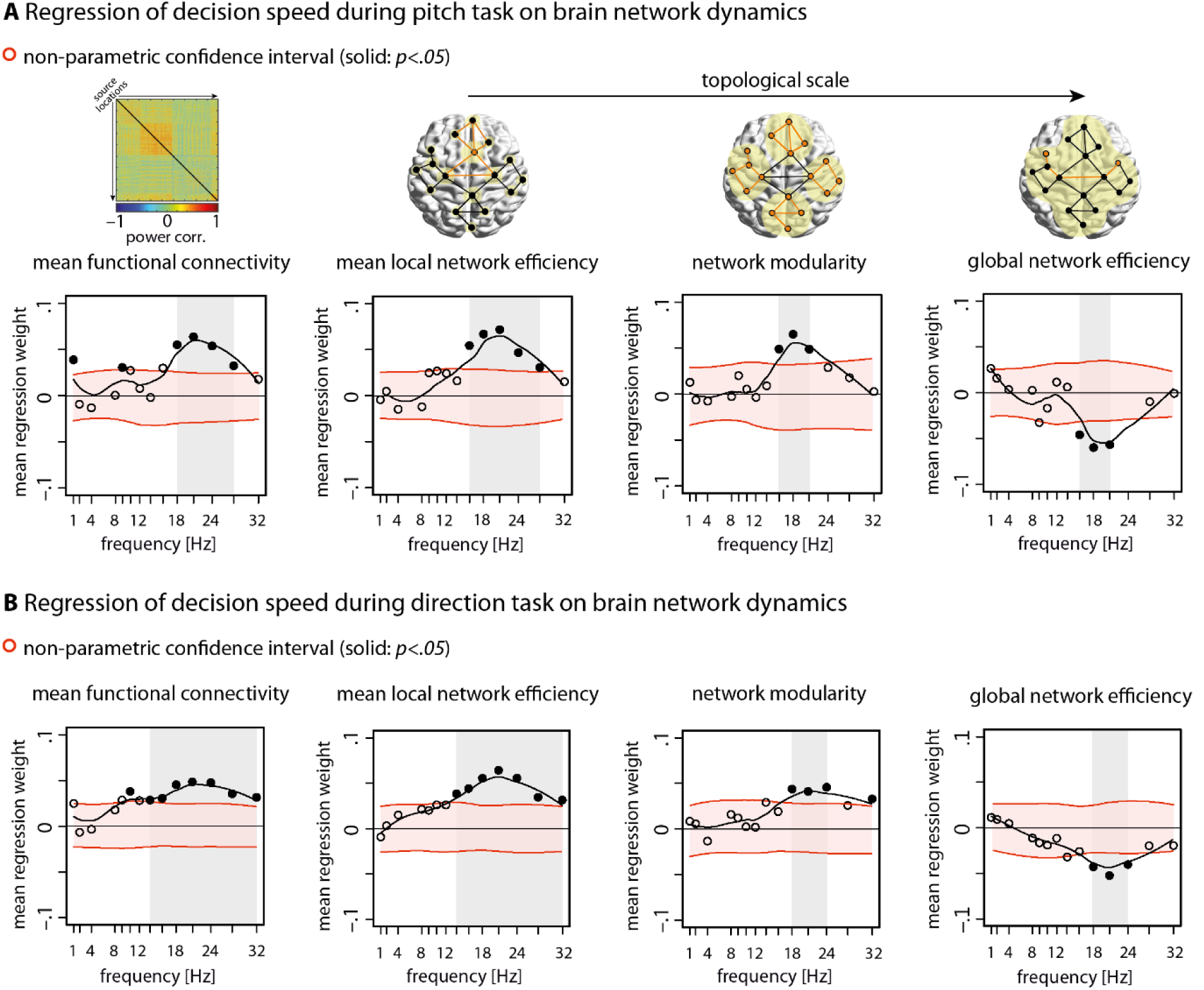
Whole-brain network dynamics of beta-band oscillations predict decision speed. The relation between the ongoing dynamics of large-scale brain networks and perceptual decisions on the auditory stimuli was investigated by means of linear regression where trial-by-trial decision speed was predicted by the temporal graph-theoretical network metrics. This analysis was done separately for each **(A)** pitch and **(B)** direction task, at frequencies ranging from 1 to 32 Hz. In the course of both auditory tasks, faster perceptual decisions positively correlated with the ongoing local efficiency (second column) and modular segregation (third column) of brain networks built upon beta-band oscillations. However, higher global integration showed the opposite effect (last column). *Black circles.* The mean regression weights averaged over subjects at each frequency. *Horizontal shades.* 95% confidence interval of the null mean regression weights generated by circularly shifting the behavioral responses across trials (corrected for multiple comparisons across frequency bins using false-coverage statement rate (FCR); p<0.05). For visualization purposes, the smooth curves were estimated using local polynomial regression fitting. *Toy graphs.* The temporal graph-theoretical metrics captured brain network states on local (local efficiency), intermediate (modularity) and global (global efficiency) scales of network topology. The yellow shaded ovals illustrate the topological scale at which a network metric is measured.

More specifically, we found positive correlations between mean functional connectivity of dynamic brain networks and decision speed in both pitch and direction task within the frequency range of 16 to 28 Hz (Figure 4, first column). In both tasks, faster perceptual decisions about the acoustic textures were accompanied by increase in mean functional connectivity of dynamic brain networks over trials. This effect was not limited to functional connectivity, and was also reflected in the topology of dynamic brain networks. On the local scale of network topology, higher mean local efficiency of dynamic brain networks within the frequency range of 16 to 28 Hz predicted faster decisions in both tasks over trials (Figure 4, second column). Moreover, on the intermediate level of network topology, higher modularity of dynamic brain networks at the same frequencies predicted faster decisions in both tasks (Figure 4, third column). Finally, on the global scale of network topology, faster decisions were predicted by decrease in global network efficiency at frequencies ranging from 18 to 21 Hz (Figure 4, second column). Note that we did not observe any significant correlation between trial-by-trial estimates of decision accuracy and brain network metrics in neither of the tasks.

The analysis of neurobehavioral correlations was based on estimating all-to-all source connectivity per trial, which covered the time points from −0.5 to 1.5 s in step of 0.05 s. To further investigate the possible prediction from pre- or post-stimulus dynamic network states, we applied the same analyses to the data measured during the pre-stimulus interval (−0.85 to 0 s) or post-stimulus interval (0 to +1 s) separately. In addition, to examine the extent to which our results might merely reflect neural processes involved in giving manual responses after the response prompt (see Figure 3), we also analyzed mid-beta band (16−28 Hz) power correlations using only the data measured during the response window (+1 s to +1.5 s). None of these analyses revealed consistent significant correlations with the speed of auditory perceptual decisions (Supplementary Figure S1).

To test the task-specificity of the correlations between a given network diagnostic and trial-by-trial decision accuracy or speed, we also computed the mean difference of the regression weights. We found no significant difference between the two tasks in predicting decision accuracy or speed from the ongoing dynamics of network topology on the whole-brain level. Finally, to investigate the possible lead/lag relationship between brain network states and trial-by-trial decision-making performance, we computed the cross-correlation between behavioral time series on the one hand and the dynamics of brain networks on the other hand. This analysis replicated the significant neurobehavioral correlations which peaked at zero trial lag (Supplementary Figure S2).

We also considered the possible effect of graph thresholding at 10% of network density on dynamic functional connectivity. To this aim, we derived the power-envelope coupling strength without thresholding the temporal graphs, and subsequently used raw trial-by-trial measures of functional connectivity in the linear regression analysis. The results were consistent with our main finding: faster perceptual decisions were positively correlated with power-envelope coupling between beta-band neural oscillations (Supplementary Figure S4). This finding suggests that functional connectivity dynamics of beta-band oscillations are not diminished by fixing the connection density of the temporal brain graphs at 10 %. Moreover, our results were also present when brain graphs were thresholded at 5% of network density (Supplementary Figure S8). Finally, In order to dissociate network from power effects, we implemented a linear regression analysis by adding the trial-by-trial estimates of source power as an additional regressor in the model. This analysis revealed that network dynamics of beta-band oscillations predicted trial-by-trial decision speed over and above the oscillatory source power (Supplementary Figure S5).

In an additional analysis, we investigated the dependence of trial-by-trial network metrics on the trial-by-trial acoustic features (i.e., spectral center and stimulus coherence) by means of separate linear regression models (consistent with the main analysis). In each model, we treated the trial-by-trial acoustic features as the dependent variable, and tested the significance of the mean regression weights averaged over subjects. On the whole-brain level, we did not observe any consistent significant correlation between brain network metrics and acoustic features in neither of the tasks (Supplementary Figure S7). In addition, our main finding - the brain-behavior relation observed on the whole-brain level - was still present when we did not control for the acoustic features of the trial-by-trial stimuli in our regression model. These together suggest that large-scale network organization of coupled beta-band oscillations during auditory perceptual decision-making is not globally altered by the external perturbation induced by the stimuli. The global configuration of brain networks is rather organized according to the decision goal based on which the auditory stimuli need to be evaluated. Overall, our findings show that the dynamics of brain functional connectivity predict trial-by-trial fluctuations in the speed at which auditory perceptual decisions are made and executed. More importantly, faster decisions positively correlated with the ongoing local clustering and modular segregation of large-scale brain networks over trials. At the same time, faster decisions were also predictable from a decrease in the global integration of dynamic brain networks. The brain network correlates of auditory perceptual decision-making were found only for decision speed, and were similar across both task sets. Additionally, our findings were specific to the mid-beta band (~20 Hz) of neural oscillations, and only observed when the neural oscillatory responses within both pre- and post-stimulus intervals were used to estimate trial-by-trial brain network states.

### Regional network states of beta-band oscillations predict decision speed

The participants judged identical acoustic stimuli under two distinct task sets. Therefore, we expected not only similar network states to correlate with auditory perceptual decision-making in both tasks (mainly associated with fronto-temporal cortical communication), but also we anticipated task-specific network states (potentially emerging from auditory association or higher-order decision areas). On the whole-brain level, we found no significant difference between the two tasks in predicting trial-by-trial speed or accuracy of the auditory perceptual decisions from the ongoing dynamics of brain networks.

However, regional properties of large-scale brain networks could still predict decision speed specifically in either pitch or direction task, or in both but in different directions. Thus, we aimed at investigating the regional network states which would differentially predict the speed of auditory perceptual decisions during the pitch versus the direction task.

Figure 5 gives a comprehensive overview of all differential network effects found at the regional level of large-scale brain networks. These maps show significant differential correlations at cortical source locations. Four regional network properties were analyzed (see Supplemental Information): (A) nodal connectivity (also known as nodal strength), (B) local efficiency, (C) modular segregation (also known as within-module z score), and (D) nodal efficiency.

**Figure 5.**
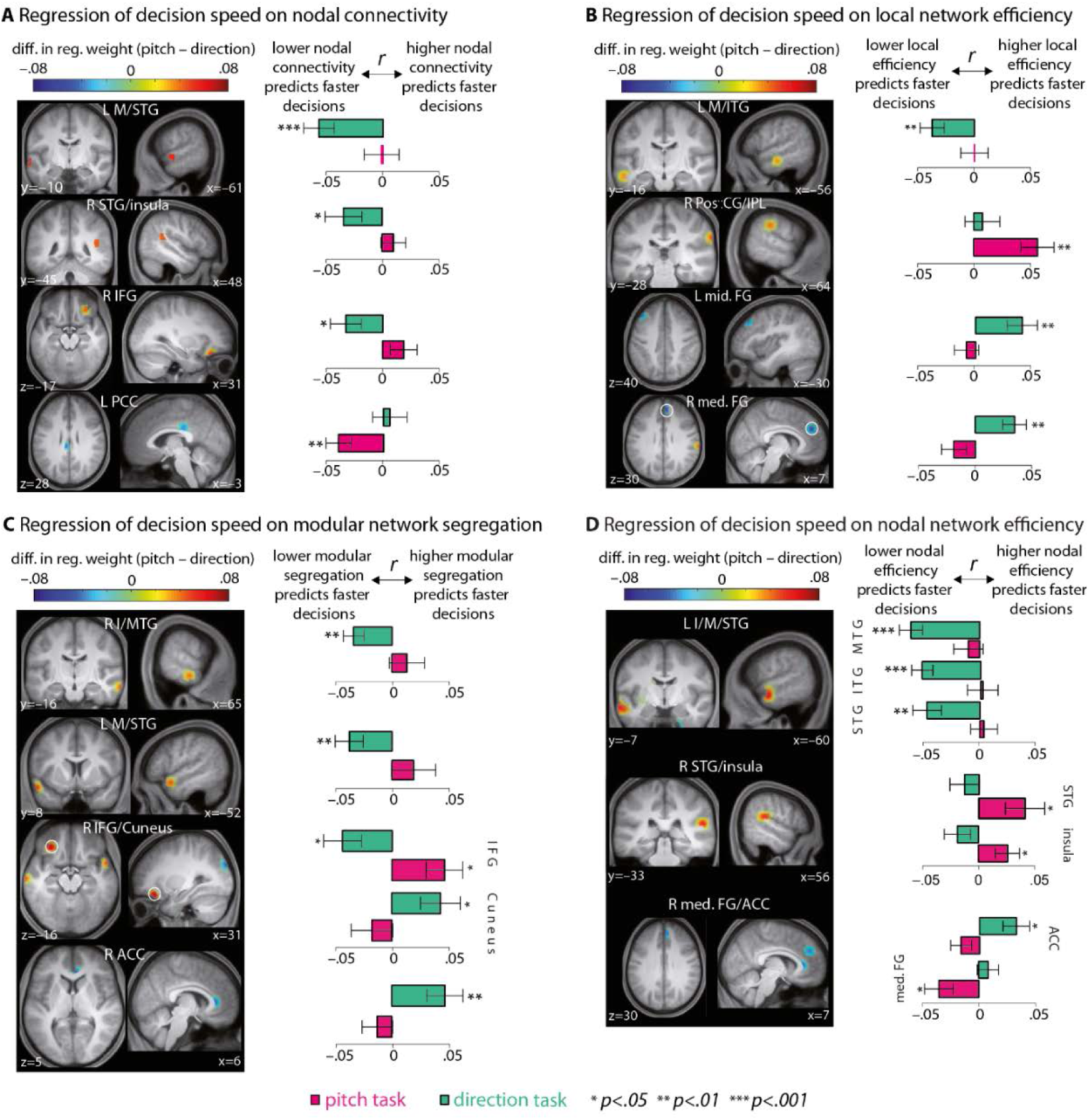
Cortical regions where network states of beta-band oscillations differentially predict decision speed during the pitch as compared to the direction task. At the regional level of large-scale brain networks, we aimed at finding task-specific correlations between the ongoing dynamics of regional network metrics and trial-by-trial decision speed. The analysis was focused on four regional network properties: **(A)** nodal connectivity, **(B)** local efficiency, **(C)** modular segregation (within-module z score) and **(D)** nodal efficiency. The direction task (green bars), as compared to the pitch task (rose bars), showed stronger correlations with network properties of the sources within temporal and frontal cortices. Within the auditory cortex, decrease in local network efficiency (B, first row), modular segregation (C, first two rows) and nodal efficiency (D, first row) supported faster decisions during the direction task. Within the frontal cortex, increase in local efficiency (B, last two rows) and module segregation (C, last row) correlated with faster decisions during the direction task. *Color maps.* For each network metric and source location, the mean regression weight obtained from the direction task data was subtracted from the mean regression weight obtained from the pitch task data. The difference was considered significant if it did not cover 95% confidence interval of the null distribution generated from shifting the behavioral responses (corrected for the number of source locations using false coverage-statement rate (FCR); p<0.05). *Bar plots.* Task-specific mean regression weights (r) whose significance was tested against zero by means of one sample permutation test with 10,000 repetitions (error bars: SEM). *Anatomical labels.* I/M/STG: inferior/middle/superior temporal gyrus; A/PCC: anterior/posterior cingulate cortex; I/mid./med. FG: inferior/middle/medial frontal gyrus; IPL: inferior parietal lobule. L and R abbreviate the left and right brain hemispheres respectively. [x,y,z] indicate MNI coordinates (mm).

First, *connectivity* of two network nodes (i.e. MEG source locations) located within left and right temporal gyri showed significant differential correlations with decision speed during the pitch task in contrast to the direction task (Figure 5A, first and second row). Lower connectivity at these locations - overlapping with middle and superior divisions of left and right temporal cortices, respectively - predicted faster auditory perceptual decisions specifically during the direction task. Also, lower connectivity of a network node in right inferior frontal gyrus predicted faster decisions during the direction task (Figure 5A, third row), whereas lower connectivity of a node overlapping with left posterior cingulate cortex predicted faster perceptual decisions during the pitch task (Figure 5A, last row).

Second, lower *local efficiency* of left middle/inferior temporal gyri specifically predicted faster decisions during the direction task (Figure 5B, first row). These faster decisions, however, were concurrent with increase in local efficiency of left middle and right medial frontal cortices (Figure 5B, last two rows).

Third, *modular segregation* of certain network nodes within left and right auditory and frontal cortices showed correlations with the decision speed specifically during the direction task (Figure 5C). More precisely, when subjects judged the overall direction of the FM tone sweeps faster, two source locations within bilateral auditory cortices showed decrease in their modular segregation (Figure 5C, first two rows). Notably, higher modular segregation of a source location within right anterior cingulate predicted faster decisions during the direction task (Figure 5C; last row).

Finally, we found strong correlations between decision speed during the direction task and integration of three nodes within left auditory cortex into the whole-brain network (Figure 5D). This result emerged from lower *nodal efficiency* of three source locations overlapping with left inferior, middle and superior temporal gyri, and was specific to the direction task (p<0.05). The correlation between decision speed during the direction task and nodal efficiency of brain networks was not limited to the regions within the auditory cortex: Higher nodal efficiency of a source location in right anterior cingulate cortex also predicted faster decisions during the direction task (Figure 5D, last row). In contrast, faster perceptual decisions during the pitch task correlated with higher nodal efficiency of right superior temporal gyrus and insula (Figure 5D, second row).

We also investigated the contribution of regional network states to the results observed on the whole-brain level shown in Figure 4. To this end, we implemented the same analysis as it was done on the whole-brain level, but used a regional network metric extracted per source location over trials as a predictor. This analysis was conducted independently per pitch and direction task. We found significant correlations between the ongoing dynamics of brain regional network metrics and trial-by-trial speed of auditory perceptual decisions in both pitch and direction tasks (non-differential effects; Supplementary Figure S3).

These results were in good agreement with the direction of the correlations observed on the whole-brain level (Figure 4). In brief, higher local network efficiency, modular segregation and nodal network efficiency of the source locations predominantly within bilateral sensorimotor and parietal cortices predicted faster decisions in both tasks. However, decrease in all of the regional network metrics in source locations predominantly within auditory cortex supported faster decisions, which was more evident in the case of the direction task.

Taken together, the results obtained at the regional level of whole-brain networks point to stronger correlations between brain network states and the speed of perceptual decisions during the direction task as compared to the pitch task. These predictions emerged from auditory and frontal cortices. Among these predictions, the stronger ones (significance level of p<0.01) converged towards decrease in nodal connectivity, local network efficiency, modular segregation and nodal network efficiency of regions within the auditory cortex. Within the frontal cortex, however, faster decisions during the direction task were predicted by increase in local network efficiency and modular segregation.

## Discussion

Time- and frequency-resolved analysis of large-scale brain networks during auditory perceptual decision-making unveiled two main results: For both pitch and direction task and on the whole-brain level, faster decisions were predicted by higher local efficiency and modular segregation, but lower global integration of coupled beta-band oscillations. On the regional level, the results of our task-differential analysis revealed that the relatively more difficult direction task relied critically on specific network configurations of temporal and frontal regions. We discuss these results in terms of neural oscillations and complex brain networks. Further elaboration is provided in Supplementary Discussion.

### Network dynamics of beta-band oscillations predict decision speed

Oscillations are key to neural communication (Adrian, 1944; Buzsaki and Draguhn, 2004; Schroeder and Lakatos, 2009; Fries, 2015). While most studies on neural oscillations aim to uncover mechanisms for dynamic excitation, inhibition, and synchrony (Singer and Gray, 1995;Womelsdorf et al., 2007;Jensen and Mazaheri, 2010; Engel et al., 2016), fewer studies have focused on long-range synchronizations between distributed cortical areas (e.g.Varela et al., 2001; Doesburg et al., 2009; Donner and Siegel, 2011; Hanslmayr et al., 2016). Here, we measured the coupling between power envelopes of MEG source signals which has been shown to underlie global network communication across cortex (Siegel et al., 2012). The temporal network dynamics of these large-scale interactions predicted the speed of auditory perceptual decisions, which was specific to brain networks tuned at mid-beta band of neural oscillations, centered around 20 Hz.

Beta-band oscillations have classically been associated with sensorimotor functions (Crone et al., 1998; Brovelli et al., 2004; Aumann and Prut, 2015) and are attenuated during voluntary movements or motor imagery (Pfurtscheller and Lopes da Silva, 1999; Pfurtscheller, 2001; Turella et al., 2016). In our study, faster decisions negatively correlated with the power of neural oscillations within alpha and beta band. However, the network effect was specific to mid-beta band. We argue that this network effect is a manifestation of large-scale neural couplings underlying auditory processing and manual responding. The results of our control analysis suggest that motor actions per se cannot explain all of the correlations we found on the whole-brain level. In this analysis, we used the mid-beta band data within the period when subjects manually reported their judgment. We observed positive correlations between decision speed and only local network efficiency (Figure S1). Knowing that local network efficiency is related to nodal clustering (Rubinov and Sporns, 2010), this effect might be due to local beta-band desynchronization coherently occurring within sensorimotor cortex, thereby forming dense clusters with high local efficiency. Beside our control analysis, we also investigated pre- and post-stimulus effects (Figure S1) which did not reveal consistent significant effects. These together highlight one very important question: What neural dynamics account for predicting decision speed from brain network states?

We here used the correlation between band-limited power envelopes as a functional connectivity measure, and had to choose a certain length for the trial-wise time windows, which is a key parameter in dynamic network analysis (Hutchison et al., 2013) and is related to the frequency content of the underlying signal. Perhaps the length of the above-mentioned time windows was not long enough to estimate correlations between beta-band power envelopes per trial. Power envelopes evolve within longer time windows as compared to their underlying carrier frequency (Siegel et al., 2012). Beta-band power envelopes fluctuate slowly at frequencies below 0.3 Hz (Engel et al., 2013) and therefore their dynamic coupling is better estimated when a time window of ~3 s (in our case one trial) is used. Accordingly, our control analysis cannot entirely preclude the effect of sensorimotor beta-band desynchronzation in our main findings. Within the last 500-ms of a trial concurrent with planning and executing a manual response, the power envelope of a ~20 Hz oscillation can be moderately modulated due to the suppression in its underlying carrier. Thus, the trial-wise power envelope correlations likely reflect the neural couplings underlying perceptual decision-making (−0.5 to +1s) and the neural underpinning of manual responses given after the response prompt (+1 to 1.5s). Indeed, the effects we found on the regional level support the involvement of auditory, sensorimotor and frontal cortices in predicting decision speed (Figure S2). However, the sluggish dynamics of beta-band power envelopes makes it difficult to dissociate network states arising from perceptual decision processes from those of manual responses.

Moreover, previous work on the timing of perceptual decision-making suggests that, in sensory-motor tasks, a decision is already represented in motor areas before a behavioral response is generated (Gold and Shadlen, 2007; de Lange et al., 2013; Kelly and O'Connell, 2015). Specifically, a decision variable undergoes a dynamic active process through which the accumulated sensory evidence is integrated over time until the action is executed (Schroeder et al., 2010; Wyart et al., 2012). Several studies across different sensory modalities have associated modulations in beta-band activity with the temporal evolution of perceptual decisions (Senkowski et al., 2006). For example, Donner et al. (2007 and 2009) reported a fronto-parietal beta-band activity which was predictive of accuracy during a visual motion detection task, and only occurred during the decision period of the trials. Accordingly, in Siegel et al. (2011) the authors provided two possible interpretations for these observations: the maintenance and accumulation of sensory evidence during the decision formation, or the maintenance of the sensorimotor mapping rule between accumulated sensory evidence and action (see Engel and Fries (2010) for an elaboration). Additionally, a study by O'Connell et al. (2012) demonstrates that, during target detection tasks in different sensory modalities, left-hemisphere beta power was modulated by a reduction in the stimulus contrast, and this gradual modulation predicted trial-by-trial reaction times. Lastly, the role of beta-band oscillations in decision-making is supported by animal studies (Heekeren et al., 2008; Haegens et al., 2011) and computational modeling (Mostert et al., 2015; Sherman et al., 2016).

In sum, and in good agreement with previous accounts (Donner and Siegel, 2011; Hipp et al., 2011), we found that large-scale network interactions mediated by the power of beta-band oscillations are crucial for perceptual decision-making. Our findings draw a direct link between the dynamic network organization of coupled neural oscillations at ~20 Hz and trial-by-trial speed of auditory perceptual decisions built up from early perception to executing manual responses.

### Network states of fronto-temporal regions supporting auditory perceptual decisions

Our task-differential analyses at the regional level suggest that the arguably more difficult direction task versus the pitch task relied critically on specific network configurations of beta-band oscillations. The differences we found in nodal network topology across the two tasks were specific to MEG sources located within temporal and frontal cortices. Within the vicinity of auditory cortex, effects in support of faster decisions during the direction task converged towards a *decrease* in (i) nodal connectivity, (ii) local efficiency, (iii) modular segregation, and (iv) nodal efficiency of source locations mostly overlapping with the anterior division of left superior temporal cortex. Within frontal cortex, however, *increase* in (i) local efficiency of left middle and medial frontal gyri, and (ii) modular segregation of right anterior cingulate, predicted faster decisions during the direction task.

One pattern forged of these results is that faster decisions during a particularly challenging auditory perceptual task are accompanied by an increase in network segregation of frontal regions, and that this segregation supports higher-order decision-related processes. Theoretically, high local clustering of neighbor nodes is associated with high efficiency in local information transfer and fault tolerance (Achard and Bullmore, 2007), indicating how well neighbor nodes can still communicate when the target node (in our case a frontal region) is removed. As such, when making a perceptual decision is relatively difficult (i.e. the direction task), the decision process benefits from a more autonomous network configuration of frontal regions.

The critical involvement of frontal regions in perceptual decision-making is supported by previous animal studies on local field potentials (LFP) where frontal cortex responses were found to selectively encode auditory stimulus features (Fritz et al., 2010) or to show higher synchrony at beta band (19-40 Hz) as the cortical representation of task rules (Buschman et al., 2012; Antzoulatos and Miller, 2016). Additionally, in the visual domain it has been shown that top-down control of attention is mediated by higher coherence between frontal and parietal LFPs at beta band (22-34 Hz) (Buschman and Miller, 2007). Recently, Stanley et al. (2016) found that higher local synchrony between LFPs over lateral prefrontal cortex within 16-20 Hz predicted stimulus categorization. Finally, a casual role for frontal cortex in perceptual decision-making was recently proposed by Rahnev et al. (2016).

In contrast to what we observed in frontal regions, network states concurrent with less clustered and less segregated auditory regions were found to speed up the decisions during the direction task. One possibility is that during these states brain networks were more integrated (the opposite pole of segregation). However, we did not find significant correlations between decision speed and the so-called module participation (Guimera and Amaral, 2005), a well-established nodal metric which is quantified based on inter-modular connectivity, and attributed to network integration and hubs (Sporns, 2007, 2013; van den Heuvel and Sporns, 2013). Accordingly, decrease in the clustering and segregation of auditory regions are likely due to pruning of some short-range (intra-modular) connections. This local network reconfiguration might be necessary to remove direct connections, and instead establish longer paths through intermediate critical nodes, thereby supporting the decision process.

We also note that faster decisions during the direction task were predicted by decrease in nodal efficiency of left auditory regions. This is perhaps due to emergence of longer paths between these regions and other network nodes, and pruning of long-range shortcuts. This globally less-efficient and more distributed information routing might be necessary to support the perceptual decisions under the more difficult direction task. Indeed, a study by Siegel et al. (2015) suggests that, during a sensorimotor decision task, information was not bounded to specific cortical regions, but instead was distributed across graded specialized cortical regions. In addition, and relevant to frequency-specific distributed information routing, the study by Weisz et al. (2014b) demonstrates that, during conscious perception, brain networks tuned at 17 Hz get more globally integrated through shorter communication paths.

To conclude, the present study suggests that large-scale network organization of coupled neural oscillations at ~20Hz (beta band) underlies how fast momentary auditory perceptual decisions are made and executed. Thus, global communication in brain networks during perceptual decision-making is likely implemented by neural oscillations at around 20 Hz. During auditory perceptual decision-making, this dynamic global communication appears as complex network interactions between beta band neural oscillations evolving within lower-order auditory and higher-order control areas.

## Materials and Methods

### Participants

Twenty healthy, right-handed volunteers (15 females, age range 20-32 years, mean±SD age=26.2±3.35 years) participated in the study. None of the participants reported any neurological diseases or hearing problems. Ethical approval was obtained from the local ethics committee of the University of Leipzig. All procedures were carried out with written informed consent of the participants and in accordance with the principles of the Declaration of Helsinki. Volunteers received a monetary compensation for participating in the study.

### Stimuli

Two auditory task sets were designed that used identical stimuli consisting of variable acoustic textures (see e.g. Overath et al., (2010); “cloud of tones” in Znamenskiy and Zador (2013)) of 400 ms duration (Figure 1A). Each stimulus consisted of 72 frequency modulated (FM) sine tone ramps, or sweeps, of 100 ms duration. The starting time points and frequencies of the individual sweeps were uniformly distributed across time and log frequency. Their frequency slopes spanned ±3.3 octaves per second. For manipulation of spectral center (low vs. high), the acoustic textures had one of eight spectral centers relative to a mean center frequency of 707.1 Hz. More specifically, they could deviate ±2, 1,0.5 or 0.25 semitones from the center frequency. Their spectral centers were approximately 630, 667, 687, 697, 717, 728, 749 and 794 Hz.

For the manipulation of the spectral coherence, a variable proportion (25, 50, 75 or 100%) of the sweeps were assigned the same frequency slope, i.e. they were “coherent” with one another. The rest of the sweeps had a randomly assigned slope.

### Tasks

The participants had to judge one feature of the auditory stimuli under two distinct task sets (Figure 1B). The feature was either the overall spectral center of the sweeps in the acoustic texture (the “pitch” feature: high or low), or the overall direction of the individual sweeps (the “direction” feature: up or down).

Trials started with a white fixation cross which appeared on a black background at the center of a back projection screen. After a pre-stimulus interval of one second (with no time jitter), an acoustic texture was presented. Then, the participants gave a delayed response to the auditory stimulus. This means, 0.6 s after the offset of the acoustic texture, the participants were visually prompted by the decision labels for the left and right hand response buttons (see Figure 1B). During the response window, the fixation cross was replaced by a question mark, asking for the subject's perceptual decision. The response to the previous trial was immediately followed (with no time jitter) by the presentation of the fixation cross of the subsequent trial.

To prevent systematic effects of motor preparation, mapping of the high/low and up/down labels on the two response buttons was randomized across trials. Participants had 3 s time at maximum to respond before the experiment would automatically proceed to the next trial. As in all MEG studies from Leipzig Max Planck center (e.g. Wöstmann et al. (2016)) the auditory stimuli were presented via a nonmagnetic, echo-free stimulus delivery system with almost linear frequency characteristic in the critical range of 200-4000 Hz.

On a separate session before MEG recording, an adaptive perceptual tracking was conducted with feedback so that each participant's average accuracy converged at ~70% in each task. This was done to avoid ceiling or floor effects, and to assure that the two tasks were challenging enough.

On the recording session, the participants completed eight task blocks, each consisting of 128 trials (~8min). The two auditory paradigms, namely the *pitch task* and the *direction task,* alternated from block to block, and the initial task was randomized across the participants. Each block contained two trials of each possible combination of direction (2 levels), coherence (4 levels) and spectral center (8 levels). There were two blocks at the beginning to familiarize the participants with the stimuli and the tasks. The last six blocks were used for data acquisition, resulting in a total of 384 trials per task for each subject. Between the blocks, participants were given self-paced breaks.

### Analysis of behavioral data

Performance of the subjects was measured using the average proportion of correct responses (i.e. accuracy) and decision speed (defined as [response time]^−1^). Decision speeds were calculated relative to the onset of the response prompt (see Figure 1B) and were used as a proxy for the difficulty of perceptual decision-making during each task. Trials on which no response was given were discarded from the analysis.

To estimate the dynamic pattern of decision accuracy over trials, a moving average procedure was applied to the trial-by-trial binary responses (i.e. correct: 1, incorrect: 0; rectangular window with unit height and length of four trials). The choice of the window size was guided by a recent study by Alavash et al. (2016) where the authors found the strongest dynamic coupling of hemodynamic brain networks and behavioral accuracy within time windows of ~16 s (four trials in our case). To capture trial-by-trial fluctuations in decision speed, we used inverse response times on trials where a decision (correct or incorrect) was given. The above procedures gave us time series estimates of decision accuracy or speed over trials per subject (Figure 1C).

To test the correlation between trial-by-trial estimates of decision accuracy or speed with trial-by-trial acoustic features (i.e. spectral center and %coherence) of the auditory stimuli, we used a two-level general linear model (GLM). In this model, time series estimates of decision accuracy or speed were predicted by an acoustic feature. Since positively-skewed distribution of response times can violate the normality assumption underlying the general linear model (Baayen and Milin, 2010), the dependent variable and the regressors were first rank-transformed, and then normalized (i.e., z-scored) before estimating the regression models (Cohen and Cavanagh, 2011).

### MEG data acquisition and preprocessing

MEG responses were recorded using a 306-channel Neuromag Vectorview system (Elekta, Helsinki, Finland) in an electromagnetically shielded room (Vacuumschmelze, Hanau, Germany) at a sampling rate of 1,000 Hz with a bandwidth of DC-330 Hz. Movement of each participant's head relative to the MEG sensors was monitored by means of five head-position measurement coils. The electroencephalogram from 64 scalp electrodes (Ag/Ag-Cl) was recorded but not analyzed in this study. The raw MEG data were first subjected to Maxfilter software to suppress disturbing magnetic interferences using the signal space separation method (Taulu et al., 2004). Next, the data were corrected for head movements and scanned for intervals where channels were static or flat. Subsequently, signals recorded from 204 planar gradiometer sensors at 102 locations were fed into the following steps which were implemented in Matlab (version 2015a; MathWorks, MA, USA) using Fieldtrip toolbox (Oostenveld et al., 2011) and other custom scripts.

The signals were high-pass filtered at 0.5 Hz (finite impulse response (FIR) filter, sinc window, order 6000, zero-phase filtering) and low-pass filtered at 140 Hz (FIR filter, sinc window, order 44, zero-phase filtering filtering). Next, epochs from –1 to +2 s around the onset of the acoustic textures were extracted and down-sampled to 500 Hz.

Independent component analysis was employed to exclude artifactual components containing heart, eye or muscle activity. Then, single-trial sensor data were visually inspected to exclude the trials still containing eye blinks or movements, muscle activity, static or flat intervals, signal jumps, drifts, or having a range larger than 300 pT/m (mean number of rejected trials±SEM: 29.4±4).

### Time-frequency analysis of MEG sensor data

To analyze the induced perturbation of the MEG signal power during trials, spectrotemporal estimates of the sensor signals were obtained within −0.5 to 1.5 s (relative to the onset of the acoustic textures), at frequencies ranging from 8 to 32 Hz on a logarithmic scale (Morlet's wavelets; number of cycles=6). The logarithm of the squared magnitude of the wavelet coefficients were then baseline-corrected relative to the power of the signals within −0.5 to 0 s.

### Source projection of MEG sensor data

Individual forward head models were created based on each participant's T1-weighted MRI image (3T Magnetom Trio, Siemens, Germany). The anatomical images were segmented using Freesurfer and coregistered to the MEG coordinates using MNE software (http://martinos.org/mne). The fit of approximately 200 digitized head surface points (Polhemus Fastrak 3D digitizer) to the reconstructed head surface was optimized using the iterative closest point algorithm after manual identification of anatomical landmarks (nasion, left, and right pre-auricular points). Individual segmented and coregistered anatomical images were spatially normalized to the standard stereotaxic MNI space. The inverse of these operations were applied to a 12-mm grid created in the template brain to obtain subject-specific grids in the standard space (1,781 inside-brain source locations with 12 mm distance).

To obtain the physical relation between sources and sensors for all grid points, single shell volume conduction models (Nolte, 2003) were constructed using the individual segmented anatomical images. The weakest of three dipole orientations per grid point was removed. Next, a linearly constrained minimum variance (LCMV) beamforming approach (Van Veen et al., 1997) was implemented. The spatial adaptive filters were generated by first concatenating all single-trial signals into one time series per subject, and then computing the covariance matrices using these time series. The regularization parameter was set to 7% and the singular value decomposition approach was used to estimate the dominant dipole orientation independently per grid point. Finally, the single-trial sensor data from −1 to +2 s around the onset of the acoustic textures were multiplied with the spatial filters, and the results were treated as trial-wise source-projected signals in the subsequent analyses.

### Time-frequency analysis of source-projected signals

Time-frequency representations of the source-projected signals were derived using Morlet's wavelets based on multiplication in the frequency domain. As in Hipp et al. (2012), the spectral band-width of the wavelets was set to 0.5 octaves (number of cycles=6). The center frequencies were spaced logarithmically using base 2 with exponents ranging from 3 to 5 in steps of 0.25. In addition, we included three lower frequencies at 1, 2 and 4 Hz to thoroughly investigate the neurobehavioral correlations within the range 1-32 Hz. Note that since our analysis was done at each trial (see below), to capture low-frequency oscillations per trial the time-frequency estimations at 1, 2 and 4 Hz were accomplished by mirror-symmetric extension of the source signals to the left and right.

For the main analysis, time points from −0.5 to 1.5 s (relative to the onset of the acoustic textures) in steps of 0.05 s were used to extract complex-valued spectrotemporal estimates of the source-projected signals per trial (41 data points). For the analysis of pre-stimulus interval, time points from −0.85 s to 0 (relative to the onset of the acoustic textures) were used. For the analysis of post-stimulus interval, we analyzed the time points from the onset of the acoustic textures up to +1 s, during which the participants listened to the auditory stimuli but did not manually give their responses. Additionally, since the power of beta-band oscillations is known to be related to preparation and execution of movements (Crone et al., 1998; Brovelli et al., 2004; Aumann and Prut, 2015), we performed one control analysis. That is, we analyzed the data within the response window (+1 s to +1.5 s relative to the onset of the acoustic textures) at frequencies within beta band (16-28 Hz). This analysis was aimed at investigating the possible effects arising from the neural processes involved in button press rather than auditory perceptual decision-making. The results of these analyses are summarized in the supplementary Figure S1.

### Correlation between ongoing neural oscillatory power and auditory perceptual decision-making

On the trial-by-trial basis, and in order to predict the individuals’ decision-making performance from the ongoing power of the source-projected signals over trials, we implemented a two-level GLM approach. First, at the single-subject level, time series estimates of trial-by-trial decision accuracy or speed (Figure 1C) were predicted by the baseline-corrected power of the whole-brain source-projected signals over trials, controlling for the effects of the acoustic features (i.e., spectral center and %coherence). For this analysis, the time points from −0.5 to 1.5 s (relative to the onset of the acoustic textures) in steps of 0.05 s were used to extract complex-valued spectrotemporal estimates of the source-projected signals per trial. Subsequently, for each pitch and direction task, separate GLMs were constructed. This procedure was applied to the data obtained from each participant, at each frequency of the neural oscillatory power (1–32 Hz). To account for the normality assumption underlying the general linear model (Baayen and Milin, 2010), the dependent variable and the predictors were first rank-transformed, and then normalized (i.e. z-scored) before estimating the regression models (Cohen and Cavanagh, 2011). Finally, the regression weights obtained from the fit of each GLM per task were averaged over participants at each frequency, and statistically compared with a null distribution of mean regression weights.

### Power envelope correlations and functional connectivity analysis

The power envelope of a band-limited oscillatory signal is the squared magnitude of the time-frequency signal following wavelet decomposition. To assess frequency-specific neural interactions, we computed Pearson's correlations between the log-transformed powers of all pairs of sources per trial (Figure 2). This analysis was done at each frequency within the range 1–32 Hz.

Prior to this analysis and to eliminate the trivial common co-variation in power measured from the same sources, we used the orthogonalization approach proposed by Hipp et al., (2012) prior to computing the power correlations (see Supplementary Information). This approach has been suggested and used to circumvent overestimation of instantaneous short-distance correlations, which can otherwise occur due to magnetic field propagation (Mehrkanoon et al., 2014; Siems et al., 2016).

The above procedure gave us frequency-specific *N-by-N* functional connectivity matrices *(N* denotes number of source locations) per subject and trial, for each pitch and direction task (Figure 2).

### Building dynamic brain networks

To construct brain graphs from functional connectivity matrices, different approaches have been suggested and used (van Wijk et al., 2010; Fornito et al., 2013; Garrison et al., 2015). One way is to construct brain graphs over different network densities by including links in the graph according to the rank of their absolute correlation values (Alexander-Bloch et al., 2010; Ginestet et al., 2011). In our study, and in order to make the brain graphs comparable in terms of size across subjects and trials, the number of links in each brain graph per trial was fixed at 10% of network density. The choice of the density threshold was based on previous work demonstrating that the brain network correlates of behavior are observed within a low-density range of network connections (Achard and Bullmore, 2007; Giessing et al., 2013; Alavash et al., 2015; Godwin et al., 2015; Alavash et al., 2016). However, to assure that the results are not specific to one network density, we repeated our analysis at 5% of network density, and summarized the results in Supplemental Figure S8.

Subsequently, binary undirected brain graphs were built, from which graph-theoretical network metrics were extracted per trial (Figure 2). The mean functional connectivity was estimated as the mean of the upper-diagonal correlation values within the sparse temporal connectivity matrices.

### Network diagnostics

Three key topological properties were estimated per trial: mean local efficiency, network modularity, and global network efficiency. These graph-theoretical metrics were used to capture dynamic patterns of functional integration and segregation on the local, intermediate, and global scales of network topology respectively (Figure 2). For each graph-theoretical metric, we computed a global network diagnostic which collapses the metric into a single measure on the whole-brain level, and a regional diagnostic characterizing the same metric but for a certain cortical source location (see Supplementary Information).

### Correlation between brain network dynamics and auditory perceptual decision-making

To predict trial-by-trial perceptual decision-making performance from the ongoing brain network states, we employed the same regression approach as we used for predicting the performance from the brain ongoing oscillatory power. That is, we implemented a GLM where time series estimates of trial-by-trial decision accuracy or speed (Figure 1C) were predicted by the graph-theoretical network metrics over trials (Figure 2), controlling for the effects of the acoustic features (i.e. spectral center and %coherence). For each network diagnostic, a separate GLM was constructed. Thus, each model consisted of three regressors, together with a constant term. Since the statistical distribution of the temporal brain network metrics is not necessarily normal, the dependent variable and the predictors were first rank-transformed, and then normalized (i.e. z-scored) before estimating the regression model (Cohen and Cavanagh, 2011). This procedure was separately applied to the data obtained from each pitch and direction task, subject, and each frequency of the neural oscillatory power. Finally, the normalized regression weights obtained from the fit of each GLM per task were averaged over participants at each frequency, and statistically compared with a null distribution of mean regression weights.

To measure the task-specificity of the correlations between a given network diagnostic and trial-by-trial decision accuracy or speed, we also computed the mean difference of the regression weights *β* obtained from each GLM model, i.e. *mean*(*β_pitch_ - β _direction_*).

### Statistical analysis

#### Behavioral data

Mean decision speeds and average accuracies were compared between the different task and stimulus conditions by means of an analysis of variance (ANOVA) for repeated measures, using “task” (pitch, direction), “coherence” (4 levels), and “spectral center” (4 levels) as within-subject factors. We used generalized eta squared (*η_G_*^2^ Bakeman (2005)) as the effect size statistic. Prior to the ANOVA, the distributions of the behavioral measures across participants were statistically analyzed by means of bootstrap Kolmogorov-Smirnov test with 10,000 repetitions to ensure that the data derive from a normally distributed population. The bootstrap Kolmogorov-Smirnov test, unlike the traditional Kolmogorov-Smirnov test, allows the presence of ties in the data (Sekhon, 2007). The behavioral measures (i.e. mean decision speed and average accuracies) were compared between the pitch and direction task using exact permutation tests for paired samples. The correlations between each of the behavioral measures across the two tasks were tested using rank-based non-parametric Spearman's *p* correlation (Spearman, 1904) with 10,000 permutations applied to the correlation coefficients (Pesarin and Salmaso, 2010). To test the correlation between trial-by-trial estimates of decision accuracy or speed with trial-by-trial acoustic features of the stimuli, a two-level GLM was separately applied to each pitch and direction task per subject. The regression weights obtained from the fit of each GLM were averaged over participants, and statistically tested against zero using one sample exact permutation tests.

#### Neurobehavioral correlations

To predict trial-by-trial perceptual decision accuracy or speed from the ongoing brain oscillatory power or network states, we implemented a two-level GLM approach. The regression weights obtained from the fit of each GLM were averaged over participants at each frequency, and statistically compared with a null distribution. The null distribution was generated using a randomization procedure where trial-by-trial binary responses or decision speed were circularly shifted 350 times (number of trials remained after preprocessing in every subject) over trials, per task and per subject. Circular shifting preserves the autocorrelation structure inherent to the time series (e.g. the trial-by-trial sequential correlation in response times (Baayen and Milin, 2010)), and thus is advantageous over random shuffling. For the circularly shifted behavioral responses, we conducted the same analysis steps as it was done for the empirical behavioral data. To statistically test the significance of the neurobehavioral correlations, the observed mean regression weight was compared with the null distribution generated from the randomization procedure at each frequency. The observed mean regression weight was considered significant if it was higher than 97.5^th^ percentile or lower than 2.5^th^ percentile of the null distribution (upper or lower bounds of the horizontal shades in Figure 3 or 4).

#### Regional analysis

At the regional level of the brain networks, it was tested whether the time series estimates of decision-making performance under each pitch and direction task differentially correlate with the regional diagnostics of brain networks (see Supplemental Information). To this end, the mean regression weights obtained from the direction task data were subtracted from the mean regression weights obtained from the pitch task data. The significance of the difference in correlations was statistically tested using a null distribution of the difference in mean regression weights generated from circularly shifting the behavioral responses. We also investigated the regional network states per pitch and direction task separately. To this end, we implemented the same analysis as it was done on the whole-brain level, but used a regional network property estimated per source location over trials. The results of this analysis is summarized in Supplementary Figure S2.

#### Significance thresholds

For all statistical tests (i.e. the inference on the behavioral and the brain network effects) we used p<0.05 (two-sided) as the threshold of significance. For the analysis on the whole-brain level, and in order to correct for multiple comparisons entailed by the number of frequency bins (14), we implemented a correction method suggested by Benjamini and Yekutieli (2005) and used in Obleser et al. (2010); Obleser and Weisz (2012). In this method, called ‘‘false coverage-statement rate’’ (FCR), we first selected those frequencies where the observed mean regression weight did not cover the null distribution at the confidence level of 95%. In a second correction pass, we (re-)constructed FCR-corrected confidence intervals for these selected frequencies at a level of 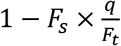, where *F_s_* is the number of selected frequencies at the first pass, *F_t_* is the total number of frequency bins tested, and *q* is the tolerated rate for false coverage statements, here 0.05. The FCR-correction procedure yields inflated, and thus more conservative confidence limits (bounds of the horizontal shades in Figure 3). To correct for multiple comparisons entailed by the regional analysis, we used the same procedure to adjust the confidence limits according to the number of source locations (1,781).

## Acknowledgements

Research was supported by the Max Planck Society (Max Planck Research Group grant to JO) and the European Research Council (ERC Consolidator grant AUDADAPT, no. 646696, to JO). Yvonne Wolff and Burkhard Maess helped acquire the MEG data.

## References

Achard S, Bullmore E (2007) Efficiency and cost of economical brain functional networks. PLoS computational biology 3:e17.

Adrian E (1944) Brain rythms. Nature 153:360–362.

Alavash M, Thiel CM, Giessing C (2016) Dynamic coupling of complex brain networks and dual-task behavior. Neuroimage 129:233–246.

Alavash M, Hilgetag CC, Thiel CM, Giessing C (2015) Persistency and flexibility of complex brain networks underlie dual-task interference. Hum Brain Mapp 36:3542–62.

Alexander-Bloch AF, Gogtay N, Meunier D, Birn R, Clasen L, Lalonde F, Lenroot R, Giedd J, Bullmore ET (2010) Disrupted modularity and local connectivity of brain functional networks in childhood-onset schizophrenia. Frontiers in systems neuroscience 4:147.

Antzoulatos EG, Miller EK (2016) Synchronous beta rhythms of frontoparietal networks support only behaviorally relevant representations. Elife 5:e17822.

Aumann TD, Prut Y (2015) Do sensorimotor beta-oscillations maintain muscle synergy representations in primary motor cortex? Trends Neurosci 38:77–85.

Baayen RH, Milin P (2010) Analyzing Reaction Times. International Journal of Psychological Research 3:12–28.

Bakeman R (2005) Recommended effect size statistics for repeated measures designs. Behavior research methods 37:379–384.

Bassett DS, Meyer-Lindenberg A, Achard S, Duke T, Bullmore E (2006) Adaptive reconfiguration of fractal small-world human brain functional networks. PNAS 103:19518–19523.

Bassett DS, Bullmore ET, Meyer-Lindenberg A, Apud JA, Weinberger DR, Coppola R (2009) Cognitive fitness of cost-efficient brain functional networks. PNAS 106:11747–11752.

Bassett DS, Wymbs NF, Rombach MP, Porter MA, Mucha PJ, Grafton ST (2013) Task-based core-periphery organization of human brain dynamics. PLoS computational biology 9:e1003171.

Benjamini Y, Yekutieli D (2005) False discovery rate-adjusted multiple confidence intervals for selected parameters. JASA 100:71–81.

Braun U, Schäfer A, Walter H, Erk S, Romanczuk-Seiferth N, Haddad L, Schweiger J, Grimm O, Heinz A, Tost H, Meyer-Lindenberg A, Bassett D (2015) Dynamic reconfiguration of frontal brain networks during executive cognition in humans. PNAS 112:11678–11683.

Brovelli A, Ding M, Ledberg A, Chen Y, Nakamura R, Bressler SL (2004) Beta oscillations in a large-scale sensorimotor cortical network: directional influences revealed by Granger causality. PNAS 101:9849–9854.

Bullmore E, Sporns O (2009) Complex brain networks: graph theoretical analysis of structural and functional systems. Nature Rev Neurosci 10:186–198.

Buschman TJ, Miller EK (2007) Top-down versus bottom-up control of attention in the prefrontal and posterior parietal cortices. Science 315:1860–1862.

Buschman TJ, Denovellis EL, Diogo C, Bullock D, Miller EK (2012) Synchronous oscillatory neural ensembles for rules in the prefrontal cortex. Neuron 76:838–846.

Buzsaki G, Draguhn A (2004) Neuronal Oscillations in Cortical Networks. Science 304:1926–1929.

Calhoun VD, Miller R, Pearlson G, Adali T (2014) The chronnectome: time-varying connectivity networks as the next frontier in fMRI data discovery. Neuron 84:262–274.

Chai LR, Mattar MG, Blank IA, Fedorenko E, Bassett DS (2016) Functional Network Dynamics of the Language System. Cereb Cortex. DOI 10.1093/cercor/bhw238.

Cohen MX, Cavanagh JF (2011) Single-trial regression elucidates the role of prefrontal theta oscillations in response conflict. Front Psychol 2:30.

Crone N, Miglioretti D, Gordon B, Lesser R (1998) Functional mapping of human sensorimotor cortex with electrocorticographic spectral analysis. II. Event-related synchronization in the gamma band. Brain 121:2301–2315.

de Lange FP, Rahnev DA, Donner TH, Lau H (2013) Prestimulus oscillatory activity over motor cortex reflects perceptual expectations. J Neuroscience 33:1400–1410.

de Pesters A, Coon WG, Brunner P, Gunduz A, Ritaccio AL, Brunet NM, de Weerd P, Roberts MJ, Oostenveld R, Fries P, Schalk G (2016) Alpha power indexes task-related networks on large and small scales: A multimodal ECoG study in humans and a non-human primate. NeuroImage 134:122–31.

Deco G, Tononi G, Boly M, Kringelbach ML (2015) Rethinking segregation and integration: contributions of whole-brain modelling. Nature Rev Neurosci 16:430–439.

Doesburg SM, Green JJ, McDonald JJ, Ward LM (2009) From local inhibition to long-range integration: a functional dissociation of alpha-band synchronization across cortical scales in visuospatial attention. Brain Res 1303:97–110.

Donner TH, Siegel M (2011) A framework for local cortical oscillation patterns. Trends in cognitive sciences 15:191–199.

Donner TH, Siegel M, Fries P, Engel AK (2009) Buildup of choice-predictive activity in human motor cortex during perceptual decision making. Curr Biol 19:1581–1585.

Donner TH, Siegel M, Oostenveld R, Fries P, Bauer M, Engel AK (2007) Population activity in the human dorsal pathway predicts the accuracy of visual motion detection. J Neurophysiol 98:345–359.

Engel AK, Fries P (2010) Beta-band oscillations--signalling the status quo? Current opinion in neurobiology 20:156–165.

Engel AK, Keitel A, Gross J (2016) Individual Human Brain Areas Can Be Identified from Their Characteristic Spectral Activation Fingerprints. PLoS Biol 14:e1002498.

Engel AK, Gerloff C, Hilgetag CC, Nolte G (2013) Intrinsic coupling modes: multiscale interactions in ongoing brain activity. Neuron 80:867–886.

Fornito A, Zalesky A, Breakspear M (2013) Graph analysis of the human connectome: promise, progress, and pitfalls. NeuroImage 80:426–444.

Fries P (2015) Rhythms for Cognition: Communication through Coherence. Neuron 88:220–235.

Friese U, Daume J, Goschl F, Konig P, Wang P, Engel AK (2016) Oscillatory brain activity during multisensory attention reflects activation, disinhibition, and cognitive control. Sci Rep 6:32775.

Fritz JB, David SV, Radtke-Schuller S, Yin P, Shamma SA (2010) Adaptive, behaviorally gated, persistent encoding of task-relevant auditory information in ferret frontal cortex. Nature Neurosci 13:1011–1019.

Garrison KA, Scheinost D, Finn ES, Shen X, Constable RT (2015) The (in)stability of functional brain network measures across thresholds. NeuroImage 118:651–661.

Giessing C, Thiel CM, Alexander-Bloch AF, Patel AX, Bullmore ET (2013) Human brain functional network changes associated with enhanced and impaired attentional task performance. J Neurosci 33:5903–5914.

Ginestet CE, Nichols TE, Bullmore ET, Simmons A (2011) Brain network analysis: separating cost from topology using cost-integration. PloS one 6:e21570.

Godwin D, Barry RL, Marois R (2015) Breakdown of the brain's functional network modularity with awareness. PNAS 112:3799–3804.

Gold JI, Shadlen MN (2007) The neural basis of decision making. Annu Rev Neurosci 30:535–574.

Guimera R, Amaral LaN (2005) Functional cartography of complex metabolic networks. Nature 433:895900.

Haegens S, Nácher V, Hernández A, Luna R, Jensen O, Romo R (2011) Beta oscillations in the monkey sensorimotor network reflect somatosensory decision making. PNAS 108:10708–10713.

Hanslmayr S, Staresina BP, Bowman H (2016) Oscillations and Episodic Memory: Addressing the Synchronization/Desynchronization Conundrum. Trends Neurosci 39:16–25.

Hanslmayr S, Gross J, Klimesch W, Shapiro KL (2011) The role of alpha oscillations in temporal attention. Brain Res Rev 67:331–343.

Heekeren HR, Marrett S, Ungerleider LG (2008) The neural systems that mediate human perceptual decision making. Nature REv Neurosci 9:467–479.

Hillebrand A, Singh KD, Holliday IE, Furlong PL, Barnes GR (2005) A new approach to neuroimaging with magnetoencephalography. Human brain mapping 25:199–211.

Hipp JF, Engel AK, Siegel M (2011) Oscillatory synchronization in large-scale cortical networks predicts perception. Neuron 69:387–396.

Hipp JF, Hawellek DJ, Corbetta M, Siegel M, Engel AK (2012) Large-scale cortical correlation structure of spontaneous oscillatory activity. Nature Neurosci 15:884–890.

Honey CJ, Kotter R, Breakspear M, Sporns O (2007) Network structure of cerebral cortex shapes functional connectivity on multiple time scales. PNAS 104:10240–10245.

Jensen O, Mazaheri A (2010) Shaping functional architecture by oscillatory alpha activity: gating by inhibition. Front Hum Neurosci 4:186.

Kayser SJ, McNair SW, Kayser C (2016) Prestimulus influences on auditory perception from sensory representations and decision processes. PNAS 113:4842–4847.

Kelly SP, O'Connell RG (2015) The neural processes underlying perceptual decision making in humans: recent progress and future directions. J Physiol Paris 109:27–37.

Kopell NJ, Gritton HJ, Whittington MA, Kramer MA (2014) Beyond the connectome: the dynome. Neuron 83:1319–1328.

Kringelbach ML, McIntosh AR, Ritter P, Jirsa VK, Deco G (2015) The Rediscovery of Slowness: Exploring the Timing of Cognition. Trends in cognitive sciences 19:616–628.

Lange J, Oostenveld R, Fries P (2013) Reduced occipital alpha power indexes enhanced excitability rather than improved visual perception. J Neurosci 33:3212–3220.

Leske S, Ruhnau P, Frey J, Lithari C, Muller N, Hartmann T, Weisz N (2015) Prestimulus Network Integration of Auditory Cortex Predisposes Near-Threshold Perception Independently of Local Excitability. Cereb Cortex 25:4898–907.

Lou B, Li Y, Philiastides MG, Sajda P (2014) Prestimulus alpha power predicts fidelity of sensory encoding in perceptual decision making. Neuroimage 87:242–251.

Marrelec G, Messe A, Giron A, Rudrauf D (2016) Functional Connectivity's Degenerate View of Brain Computation. PLoS computational biology 12:e1005031.

Mehrkanoon S, Breakspear M, Britz J, Boonstra TW (2014) Intrinsic coupling modes in source-reconstructed electroencephalography. Brain Connectivity 4:812–825.

Misic B, Sporns O (2016) From regions to connections and networks: new bridges between brain and behavior. Current opinion in neurobiology 40:1–7.

Misic B, Betzel RF, de Reus MA, van den Heuvel MP, Berman MG, McIntosh AR, Sporns O (2016) Network-Level Structure-Function Relationships in Human Neocortex. Cereb Cortex 26:3285–3296.

Mostert P, Kok P, de Lange FP (2015) Dissociating sensory from decision processes in human perceptual decision making. Sci Rep 5:18253.

Müller N, Weisz N (2012) Lateralized auditory cortical alpha band activity and interregional connectivity pattern reflect anticipation of target sounds. Cereb Cortex 22:1604–1613.

Nicol RM, Chapman SC, Vertes PE, Nathan PJ, Smith ML, Shtyrov Y, Bullmore ET (2012) Fast reconfiguration of high-frequency brain networks in response to surprising changes in auditory input. J Neurophysiol 107:1421–1430.

Nolte G (2003) The magnetic lead field theorem in the quasi-static approximation and its use for magnetoencephalography forward calculation in realistic volume conductors. Physics in medicine and biology 48.

O'Connell RG, Dockree PM, Kelly SP (2012) A supramodal accumulation-to-bound signal that determines perceptual decisions in humans. Nature neuroscience 15:1729–1735.

Obleser J, Weisz N (2012) Suppressed alpha oscillations predict intelligibility of speech and its acoustic details. Cereb Cortex 22:2466–2477.

Obleser J, Leaver AM, Vanmeter J, Rauschecker JP (2010) Segregation of vowels and consonants in human auditory cortex: evidence for distributed hierarchical organization. Front Psychol 1:232.

Oostenveld R, Fries P, Maris E, Schoffelen JM (2011) FieldTrip: Open source software for advanced analysis of MEG, EEG, and invasive electrophysiological data. Computational intelligence and neuroscience 2011:156869.

Overath T, Kumar S, Stewart L, von Kriegstein K, Cusack R, Rees A, Griffiths TD (2010) Cortical mechanisms for the segregation and representation of acoustic textures. J Neurosci 30:2070–2076.

Palva JM, Monto S, Kulashekhar S, Palva S (2010) Neuronal synchrony reveals working memory networks and predicts individual memory capacity. PNAS 107:7580–7585.

Palva S, Palva JM (2012) Discovering oscillatory interaction networks with M/EEG: challenges and breakthroughs. Trends in cognitive sciences 16:219–230.

Park H, Ince RA, Schyns PG, Thut G, Gross J (2015) Frontal top-down signals increase coupling of auditory low-frequency oscillations to continuous speech in human listeners. Current biology : CB 25:1649–1653.

Park HJ, Friston K (2013) Structural and functional brain networks: from connections to cognition. Science 342:1238411.

Pesarin F, Salmaso L (2010) The permutation testing approach: a review. Statistica 70:481–509.

Pfurtscheller G (2001) Functional brain imaging based on ERD/ERS. Vision Research 41:1257–1260.

Pfurtscheller G, Lopes da Silva FH (1999) Event-related EEG/MEG synchronization and desynchronization: basic principles. Clinical neurophysiology 110:1842–1857.

Rahnev D, Nee DE, Riddle J, Larson AS, D'Esposito M (2016) Causal evidence for frontal cortex organization for perceptual decision making. PNAS 113:6059–6064.

Rubinov M, Sporns O (2010) Complex network measures of brain connectivity: uses and interpretations. NeuroImage 52:1059–1069.

Sadaghiani S, Kleinschmidt A (2016) Brain Networks and alpha-Oscillations: Structural and Functional Foundations of Cognitive Control. Trends in cognitive sciences 20:805–817.

Sadaghiani S, Poline JB, Kleinschmidt A, D'Esposito M (2015) Ongoing dynamics in large-scale functional connectivity predict perception. PNAS 112:8463–8468.

Schroeder CE, Lakatos P (2009) Low-frequency neuronal oscillations as instruments of sensory selection. Trends Neurosci 32:9–18.

Schroeder CE, Wilson DA, Radman T, Scharfman H, Lakatos P (2010) Dynamics of Active Sensing and perceptual selection. Current opinion in neurobiology 20:172–176.

Sekhon J (2007) Multivariate and propensity score matching software with automated balance optimization: The matching package for R. J Stat Softw 42.

Senkowski D, Molholm S, Gomez-Ramirez M, Foxe JJ (2006) Oscillatory beta activity predicts response speed during a multisensory audiovisual reaction time task: a high-density electrical mapping study. Cereb Cortex 16:1556–1565.

Shen K, Hutchison RM, Bezgin G, Everling S, McIntosh AR (2015) Network structure shapes spontaneous functional connectivity dynamics. J Neurosci 35:5579–5588.

Sherman MA, Lee S, Law R, Haegens S, Thorn CA, Hamalainen MS, Moore CI, Jones SR (2016) Neural mechanisms of transient neocortical beta rhythms: Converging evidence from humans, computational modeling, monkeys, and mice. PNAS

Shine JM, Bissett PG, Bell PT, Koyejo O, Balsters JH, Gorgolewski KJ, Moodie CA, Poldrack RA (2016) The dynamics of functional brain networks: Integrated network states during cognitive task performance. Neuron 92:1–11.

Siegel M, Engel AK, Donner TH (2011) Cortical network dynamics of perceptual decision-making in the human brain. Frontiers in human neuroscience 5:21.

Siegel M, Donner TH, Engel AK (2012) Spectral fingerprints of large-scale neuronal interactions. Nature reviews Neuroscience 13:121–134.

Siegel M, Buschman TJ, Miller EK (2015) Cortical information flow during flexible sensorimotor decisions. 348.

Siems M, Pape AA, Hipp JF, Siegel M (2016) Measuring the cortical correlation structure of spontaneous oscillatory activity with EEG and MEG. NeuroImage 129:345–355.

Singer W, Gray CM (1995) Visual feature integration and the temporal correlation hypothesis. Annu Rev Neurosci 18:555–586.

Spearman C (1904) The proof and measurement of association between two things. The American Journal of Psychology 15:72–101.

Sporns O, Honey CJ, Kötter R (2007) Identification and Classification of Hubs in Brain Networks. PLoS One 17:e1049.

Sporns O (2013) Network attributes for segregation and integration in the human brain. Current opinion in neurobiology 23:162–171.

Sporns O (2014) Contributions and challenges for network models in cognitive neuroscience. Nature neuroscience 17:652–660.

Stanley DA, Roy JE, Aoi MC, Kopell NJ, Miller EK (2016) Low-Beta oscillations turn up the gain during category judgments. Cerebral Cortex.

Strauß A, Wöstmann M, Obleser J (2014) Cortical alpha oscillations as a tool for auditory selective inhibition. Front Human Neuroscie 8:350.

Taulu S, Kajola M, Simola J (2004) Suppression of interference and artifacts by the signal space separation method. Brain topography 16:269–275.

Turella L, Tucciarelli R, Oosterhof NN, Weisz N, Rumiati R, Lingnau A (2016) Beta band modulations underlie action representations for movement planning. Neuroimage 136:197–207.

van den Heuvel MP, Sporns O (2013) Network hubs in the human brain. Trends in cognitive sciences 17:683–696.

Van Veen BD, van Drongelen W, Yuchtman M, Suzuki A (1997) Localization of Brain Electrical Activity via Linearly Constrained Minimum Variance Spatial Filtering. IEEE Trans Biomed Engineer 44.

van Wijk BCM, Stam CJ, Daffertshofer A (2010) Comparing Brain Networks of Different Size and Connectivity Density Using Graph Theory. PloS one 5:e13701.

Varela F, Lachaux J, Rodriguez E, Martinerie J (2001) The brain web: Phase synchronization and large-scale integration. Nature Rev Neurosci 2:229–239.

Weisz N, Muller N, Jatzev S, Bertrand O (2014a) Oscillatory alpha modulations in right auditory regions reflect the validity of acoustic cues in an auditory spatial attention task. Cereb Cortex 24:25792590.

Weisz N, Wuhle A, Monittola G, Demarchi G, Frey J, Popov T, Braun C (2014b) Prestimulus oscillatory power and connectivity patterns predispose conscious somatosensory perception. PNAS 111:E417–425.

Womelsdorf T, Schoffelen JM, Oostenveld R, Singer W, Desimone R, Engel AK, Fries P (2007) Modulation of neuronal interactions through neuronal synchronization. Science 316:1609–1612.

Wöstmann M, Herrmann B, Maess B, Obleser J (2016) Spatiotemporal dynamics of auditory attention synchronize with speech. PNAS 113: 3873–3878.

Wyart V, de Gardelle V, Scholl J, Summerfield C (2012) Rhythmic fluctuations in evidence accumulation during decision making in the human brain. Neuron 76:847–858.

Zalesky A, Fornito A, Cocchi L, Gollo LL, Breakspear M (2014) Time-resolved resting-state brain networks. PNAS 111:10341–10346.

Znamenskiy P, Zador AM (2013) Corticostriatal neurons in auditory cortex drive decisions during auditory discrimination. Nature 497:482–485.

